# Multiplex gene editing by CRISPR-Cpf1 through autonomous processing of a single crRNA array

**DOI:** 10.1101/049122

**Authors:** Bernd Zetsche, Matthias Heidenreich, Prarthana Mohanraju, Iana Fedorova, Jeroen Kneppers, Ellen M. DeGennaro, Nerges Winblad, Sourav R. Choudhury, Omar O. Abudayyeh, Jonathan S. Gootenberg, Wen Y. Wu, David A. Scott, Konstantin Severinov, John van der Oost, Feng Zhang

## Abstract

Microbial CRISPR-Cas defense systems have been adapted as a platform for genome editing applications built around the RNA-guided effector nucleases, such as Cas9. We recently reported the characterization of Cpf1, the effector nuclease of a novel type V-A CRISPR system, and demonstrated that it can be adapted for genome editing in mammalian cells (Zetsche et al., 2015). Unlike Cas9, which utilizes a trans-activating crRNA (tracrRNA) as well as the endogenous RNaseIII for maturation of its dual crRNA:tracrRNA guides (Deltcheva et al., 2011), guide processing of the Cpf1 system proceeds in the absence of tracrRNA or other Cas (CRISPR associated) genes (Zetsche et al., 2015) (Figure 1a), suggesting that Cpf1 is sufficient for pre-crRNA maturation. This has important implications for genome editing, as it would provide a simple route to multiplex targeting. Here, we show for two Cpf1 orthologs that no other factors are required for array processing and demonstrate multiplex gene editing in mammalian cells as well as in the mouse brain by using a designed single CRISPR array.

---

To confirm our previous observation that Cpf1 alone is sufficient for array processing (Zetsche et al., 2015), we synthesized an artificial CRISPR pre-crRNA array consisting of four spacers separated by direct repeats (DRs) from the CRISPR locus of *Francisella novicida* (FnCpf1) locus and tested the pre-crRNA processing activity in vitro of two Cpf1 orthologs with activity in mammalian cells, *Acidaminococcus* Cpf1 (AsCpf1) and *Lachnospiraceae* Cpf1 (LbCpf1). Both AsCpf1 and LbCpf1 cleaved the CRISPR array in similar patterns within 10 min, whereas they did not cleave a control RNA lacking DR sequence features (Figure 1b). These findings are consistent with a recent report that FnCpf1 is the only required component for crRNA processing in vitro and in heterologous systems (Fonfara et al., 2016). The typical cleavage pattern suggests that Cpf1 recognizes secondary structures and/or motifs on its pre-crRNA array, which is also in agreement with observations from FnCpf1 (Fonfara et al., 2016). Moreover, it reinforces our previous finding that the DR sequences of Cpf1 family proteins are highly conserved and functionally interchangeable (Zetsche et al., 2015). We analyzed an array cleaved with AsCpf1 by small RNAseq and confirmed that the cleavage products correlate to fragments resulting from cuts at the 5’ end of each DR hairpin, identical to the cleavage pattern we observed previously in *E. coli* heterologously expressing FnCpf1 CRISPR systems (Zetsche et al., 2015) (Figure 1c).

**Figure 1.**
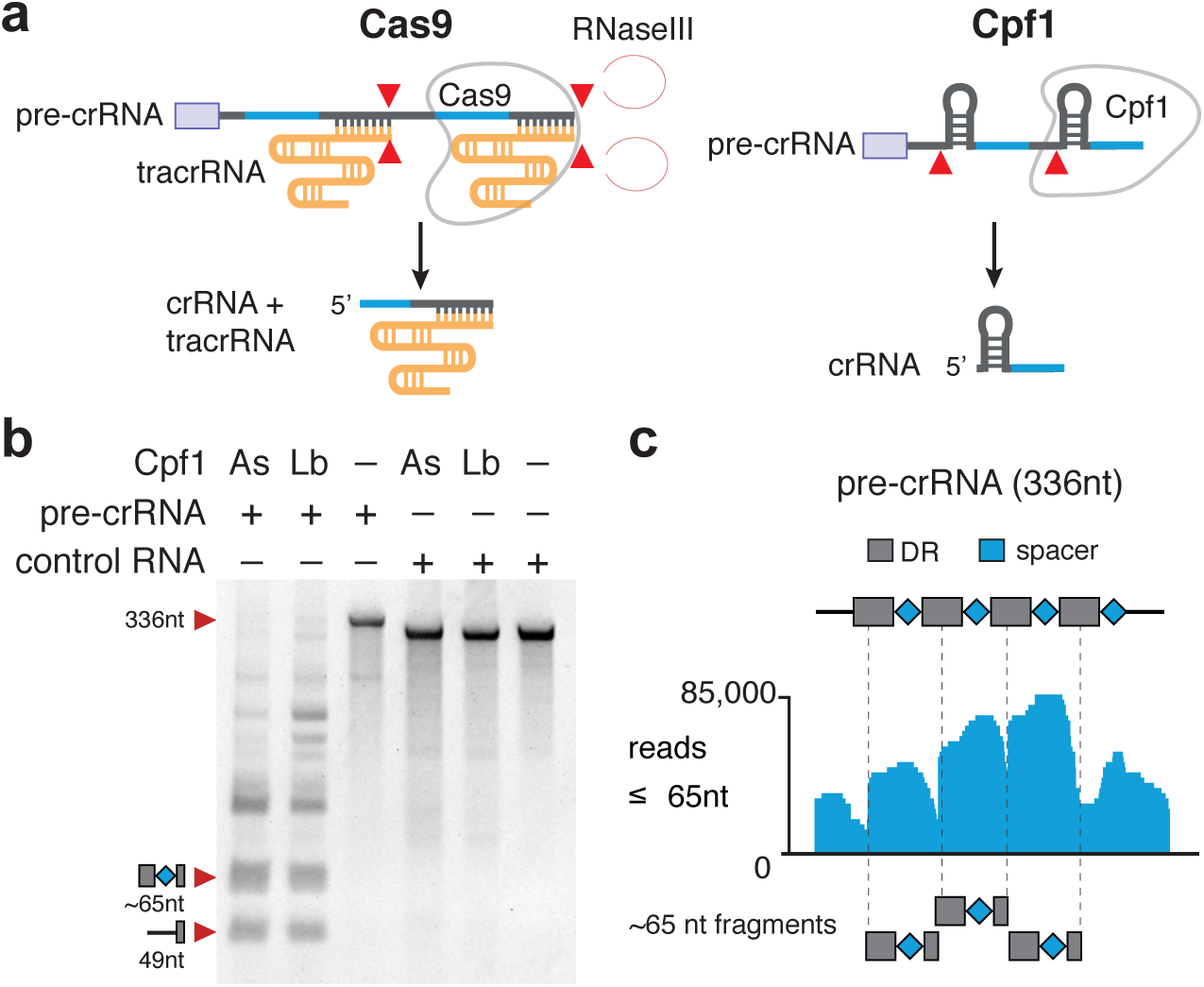
Cpf1 mediates processing of pre-crRNA. (**a**) Schematic of pre-crRNA processing for Cas9 and Cpf1. Cleavage sites indicated with red triangle. (**b**) *In vitro* processing of FnCpf1 pre-crRNA transcript (80 nM) with purified AsCpf1 or LbCpf1 protein (∼320 nM). In the presence of Cpf1 nuclease the pre-crRNA was cleaved in a distinct pattern, indicating cleavage at similar sequence motifs. RNA molecules without Cpf1 DR features where not cleaved by Cpf1 (control RNA). (**c**) RNAseq analysis of FnCpf1 pre-crRNA cleavage products, as shown in (b). A high fraction of sequence reads smaller than 65nt are cleavage products of spacers flanked by DR sequences.

We further validated these results by generating Cpf1 mutants that are unable to process arrays. Guided by the crystal structure of AsCpf1 (Yamano et al., 2016), we identified five conserved amino acid residues close to the 5’ end of the DR that are likely to disrupt array processing and mutated them to alanine (H800, K809, K860, F864, and R790) (Yamano et al., 2016). All of these mutations interfered with pre-crRNA processing but not with DNA cleavage activity *in vitro* (Figure 2a), an effect that was also observed with conserved residues in FnCpf1 (Fonfara et al., 2016). As with FnCpf1 mediated pre-crRNA processing (Fonfara et al., 2016), AsCpf1 recognizes specific nucleotides at the 5’ flank of the DR stem loop (U22-A37). Substitution at positions A16 and U17 did not affect pre-crRNA processing, substitution at position A18 and U21 weakens RNA cleavage, and A19 and U20 completely abolished RNA cleavage (Figure 2b). Although this indicates that the RNAse activity of Cpf1 is sequence specific, it would nevertheless be helpful in some genome editing applications to have DNAse activity without any RNase activity. We therefore conducted dosage tests (varying both the amount of protein and of the incubation time) on the five alanine mutants to determine if they completely abolish array processing or just reduce it. In *in vitro* reactions four of the AsCpf1 mutants (K809A, K860A, F864A, and R790A) show pre-crRNA processing when used at high concentration (Figure 2c) or for extended incubation times (Figure 2d), but H800A was inactive regardless of dose and time.

**Figure 2.**
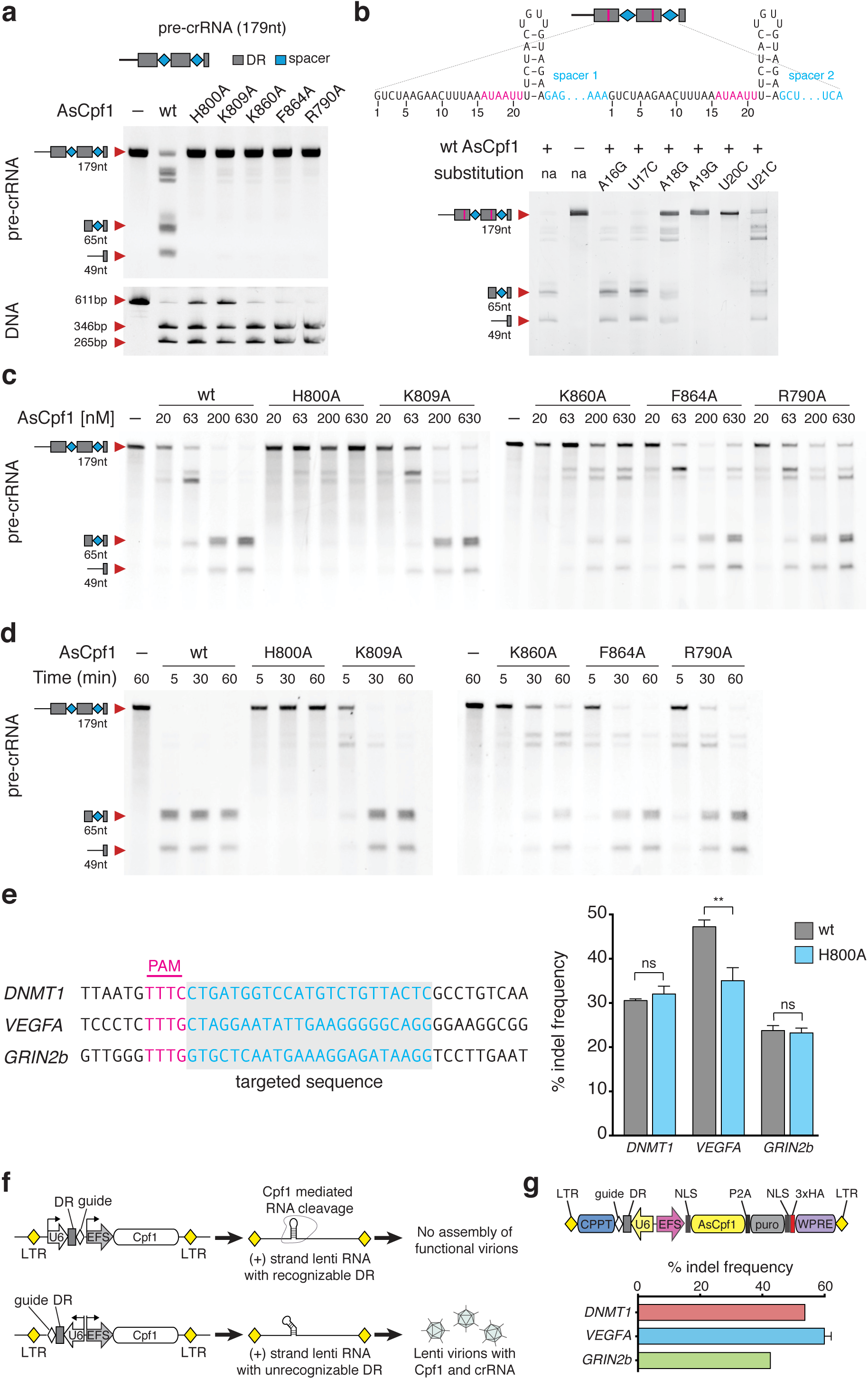
Cpf1 mediated pre-crRNA processing is independent of DNA cleavage. (**a**) Pre-crRNA (top) and DNA cleavage (bottom) mediated by AsCpf1 point mutants. H800A, K809A, K860A, F864A, and R790A fail to process pre-crRNA but retain DNA cleavage activity *in vitro*. 330 nM pre-crRNA was cleaved with 500 nM Cpf1 in 15 min and 25 nM DNA was cleaved with 165 nM Cpf1 in 30 min. (**b**) Cpf1 mediated pre-crRNA processing is sequence dependent. Single nucleotide substitutions at position A19 and U20 abolish RNA cleavage *in vitro*. 200 nM pre-crRNA was cleaved with 500 nM Cpf1 in 1 hour. (**c**, **d**) AsCpf1 point mutants, with the exception of H800A, are active at high dose. (**c**) Titration of AsCpf1 mutants reveals pre-crRNA processing at high AsCpf1 protein concentration. (**d**) Prolonged incubation time allows pre-crRNA processing by AsCpf1 point mutants. Only H800A does not process pre-crRNA to mature crRNA at high dose. 165 nM pre-crRNA was incubated with the indicated concentration (**c**) or with 500 nM AsCpf1 protein (**d**) for 30 min. (**e**) Schematic of genes targeted in HEK293T cells (left) and indels mediated by wt and H800A mutant AsCpf1 in HEK293T cells quantified using SURVEYOR nuclease assay (right). Indel frequencies mediated by H800A mutant AsCpf1 are comparable to wt AsCpf1, bars are mean of 3 technical replicates, error bars are SEM. (Student *t*-test; ns = not significant; ** = p-value 0.003) (**f**) Schematic of lenti-Cpf1 construct with the U6::DR cassette in different orientations, (+)-strand lenti RNA with recognizable DRs are susceptible for Cpf1 mediated degradation and do not result in functional virion formation (i.e. 100% cell death after puromycin treatment). (**g**) Schematic of lenti-AsCpf1 (pY108) construct (top), and indel frequencies in lenti-AsCpf1 transduced HEK cells. Indels were analyzed by SURVEYOR nuclease assay after puromycin selection 10 days after transduction. U6, Pol III promoter; CMV, cytomegalovirus promoter; NLS, nuclear localization signal; HA, hemagglutinin tag; DR, direct repeat sequence; P2A, porcine teschovirus-1 2A self-cleaving peptide; LTR, long terminal repeat; WPRE, Woodchuck Hepatitis virus posttranscriptional regulatory element.

We next tested if this mutant retains activity in human cells. We used guides targeting *DNMT1*, *VEGFA*, and *GRIN2b* (Figure 2e) to compare formation of insertion/deletion (indel) events mediated by wild-type or mutant AsCpf1 in human embryonic kidney (HEK) 293T cells. Indel frequency at the *DNMT1* and *GRIN2b* loci were identical between wild-type and H800A AsCpf1, whereas indel frequencies at the *VEGFA* locus were higher in cells transfected with wild-type AsCpf1. Taken together this data show that RNA and DNA cleavage activity can be separated in Cpf1 proteins, allowing DNA editing without potential interference with endogenous RNAs in mammalian cells.

Cpf1 mediated RNA cleavage needs to be considered when designing lenti-virus vectors for simultaneous expression of nuclease and guide (Figure 2f). Lenti virions carry a (+) strand RNA copy of the sequence flanked by long terminal repeats (LTR), allowing Cpf1 to bind and cleave at DR sequences. Hence, reversing the orientation of the DR is expected to result in (+) strand lenti RNAs not susceptible to Cpf1 mediated cleavage. We designed a lenti vector encoding AsCpf1 together with a puromycin resistance gene and a crRNA expression cassette. We transduced HEK293T cells with a MOI (multiplicity of infection) of <0.3 and analyzed indel frequencies in puromycin selected cells 10 days post infection. Using single guides encoded on a reversed expression cassette targeting *DNMT1*, *VEGFA*, or *GRIN2b* resulted in robust indel formation for each targeted gene (Figure 2g).

We leveraged the simplicity of Cpf1 crRNA maturation to achieve multiplex genome editing in HEK293T cells using customized CRISPR arrays. We chose four guides targeting different genes (*DNMT1*, *EMX1*, *VEGFA*, and *GRIN2b*) and constructed three variant arrays for expression of pre-crRNAs. In array-1, crRNAs in their mature form (19nt DR with 23nt guide) were expressed. In array-2, crRNAs are in an intermediate form (19nt DR with 30nt guide), and array-3 was designed to present crRNAs in their unprocessed form (35nt DR with 30nt guides). All arrays contain the DR from AsCpf1 and were driven by a U6 promoter from the same plasmid as AsCpf1 (Figure 3a). Indel events could be detected at each targeted locus in cells transfected with array-1 or −2. However, the crRNA targeting *EMX1* resulted in indel frequencies of <2% when expressed from array-3. Overall, array-1 performed best, with all guides showing indel levels comparable to those mediated by single crRNAs (Figure 3b). Furthermore, small RNAseq performed with RNA isolated from HEK cells transfected with array-1 and AsCpf1 confirms that autonomous, Cpf1-mediated pre-crRNA processing occurs in mammalian cells (Figure 3c). Using arrays with guides in different order result in similar indel frequencies, suggesting that positioning within an array is not crucial for activity (**Supplementary figure 1**).

**Figure 3.**
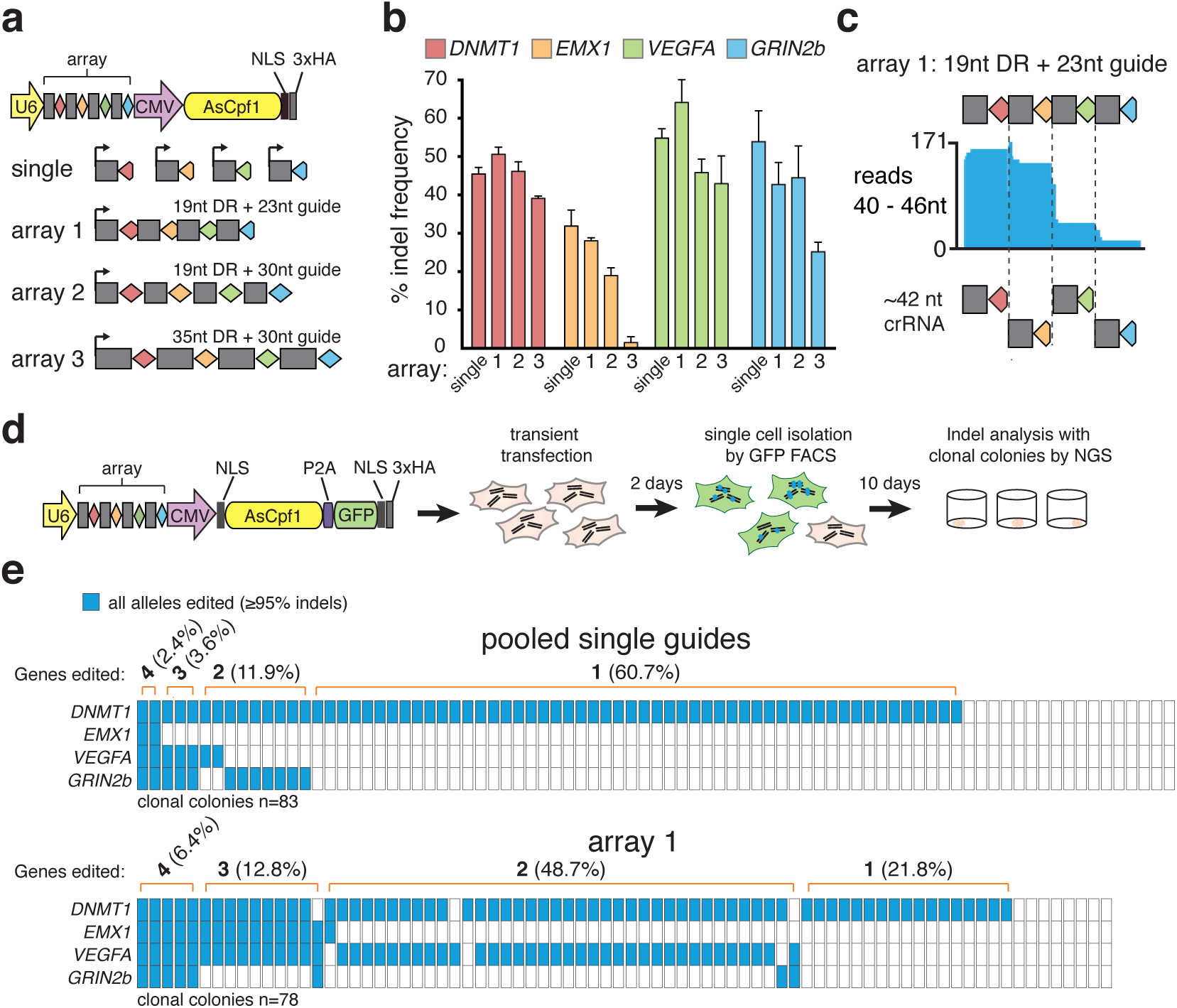
Cpf1 mediated processing of pre-crRNA transcripts allows multiplex gene editing in mammalian cells. (**a**) Schematic of multiplex gene editing with AsCpf1, using a single plasmid approach. (**b**) Genome editing at four different genomic loci mediated by AsCpf1 with different versions of artificial CRISPR arrays. Array-1 is made of 19nt DRs and 23nt guides, array 2 is made of 19nt DRs and 30nt guides and array-3 is made of 35nt DRs and 30nt guides. Indels were analyzed by SURVEYOR nuclease assay 3 days post transfection; bars are mean of two individual experiments with 3-5 technical replicates, error bars are SEM. (**c**) Small RNAseq reads from HEK cells transfected with AsCpf1 and array-1 show fragments corresponding to mature crRNA for each of the four guides. (**d**) Experimental outline for analysis of indel events in clonal colonies 48 hours after transient transfection. (**e**) Quantification of indel events measured by NGS in clonal colonies from HEK cells transient transfected with pooled single guide plasmids or plasmid carrying array-1. Colonies were expanded for 10 days after sorting. Each column represents one clonal colony, blue rectangles indicate target genes with all alleles edited.

To confirm that multiplex editing occurs within single cells, we generated AsCpf1-P2A-GFP constructs to enable fluorescence-activated cell sorting (FACS) of transduced single cells (Figure 3d) and clonal expansion. We used next generation deep sequencing (NGS) to compare edited loci within clonal colonies derived from cells transfected with either pooled single guides or array-1 construct. Focusing on targeted genes edited at every locus (indels ≥95%) shows that multiplex editing occurs more frequently in colonies transfected with array-1 (6.4% all targets, 12.8% three targets, 48.7% two targets) than in pooled transfection (2.4% all targets, 3.6% three targets, 11.9% two targets).

Multiplex genome editing provides a powerful tool for targeting members of multigene families, dissecting gene networks, and modeling multigenic disorders in vivo (Dominguez et al., 2016; Heidenreich and Zhang, 2015; Hsu et al., 2014). We therefore tested multiplex genome editing in neurons using AsCpf1. Delivery of CRISPR-effector proteins and guide RNAs into the mammalian brain is challenging, in particular due to its high complexity and the presence of the blood-brain barrier. One of the most promising delivery methods is based on viral vectors, which can be efficiently administered locally into brain parenchyma (Choudhury et al., 2016). We designed a gene-delivery system based on adeno-associated viral vectors (AAVs) for expression of AsCpf1 in neurons. We generated a dual vector system in which AsCpf1 and the CRISPR-Cpf1 array were cloned separately (Figure 4a). The human Synapsin I promoter (hSyn1) was used to restrict expression of AsCpf1 to neurons. For the Cpf1-array vector, we constructed a U6 promoter-driven Cpf1 array targeting the neuronal genes *Mecp2*, *Nlgn3*, and *Drd1*. Mutations in these genes have been implicated in mental illness including schizophrenia and autism spectrum disorders (Allen et al., 2008; Amir et al., 1999; Jamain et al., 2003). We also included an hSyn1-driven expression cassette for the green fluorescent protein (GFP) fused to the KASH nuclear transmembrane domain (Ostlund et al., 2009) to enable FACS of targeted cell nuclei from injected brain tissues (Swiech et al., 2015).

We first transduced mouse primary cortical neurons *in vitro* and observed robust expression of AsCpf1 and GFP-KASH one week after viral delivery. SURVEYOR nuclease assay on purified neuronal DNA confirmed indel formations in all three targeted genes (**Supplementary figure 2**). Next, we tested whether AsCpf1 can be expressed in the brains of living mice for multiplex genome editing *in vivo*. We stereotactically injected our dual vector system in a 1:1 ratio into the hippocampal dentate gyrus (DG) of adult male mice. Four weeks after viral delivery we observed robust expression of AsCpf1 and GFP-KASH in the DG (Figure 4b–d). Consistent with previous studies (Konermann et al., 2013; Swiech et al., 2015), we observed ∼75% co-transduction efficiency of the dual viral vectors (Figure 4c). We isolated targeted DG cell nuclei by FACS (Supplementary figure 3) and quantified indel formation using NGS. We found indels in all three targeted loci with ∼23%, ∼38%, and ∼51% indel formation in *Mecp2*, *Nlgn3*, and *Drd1*, respectively (Figure 4e,f). We next quantified the effectiveness of biallelic disruption of the autosomal gene *Drd1* and found ∼47% of all sorted nuclei (i.e. ∼87% of all Drd1-edited cells) harbored biallelic modifications (Figure 4g). Next, we quantified the multiplex targeting efficiency in single neuronal nuclei. Our results show that ∼17% of all transduced neurons were modified in all three targeted loci (Figure 4h). Taken together, our results demonstrate the effectiveness of AAV-mediated delivery of AsCpf1 into the mammalian brain and simultaneous multi-gene targeting *in vivo* using a single array transcript driven by the U6 promoter.

**Figure 4.**
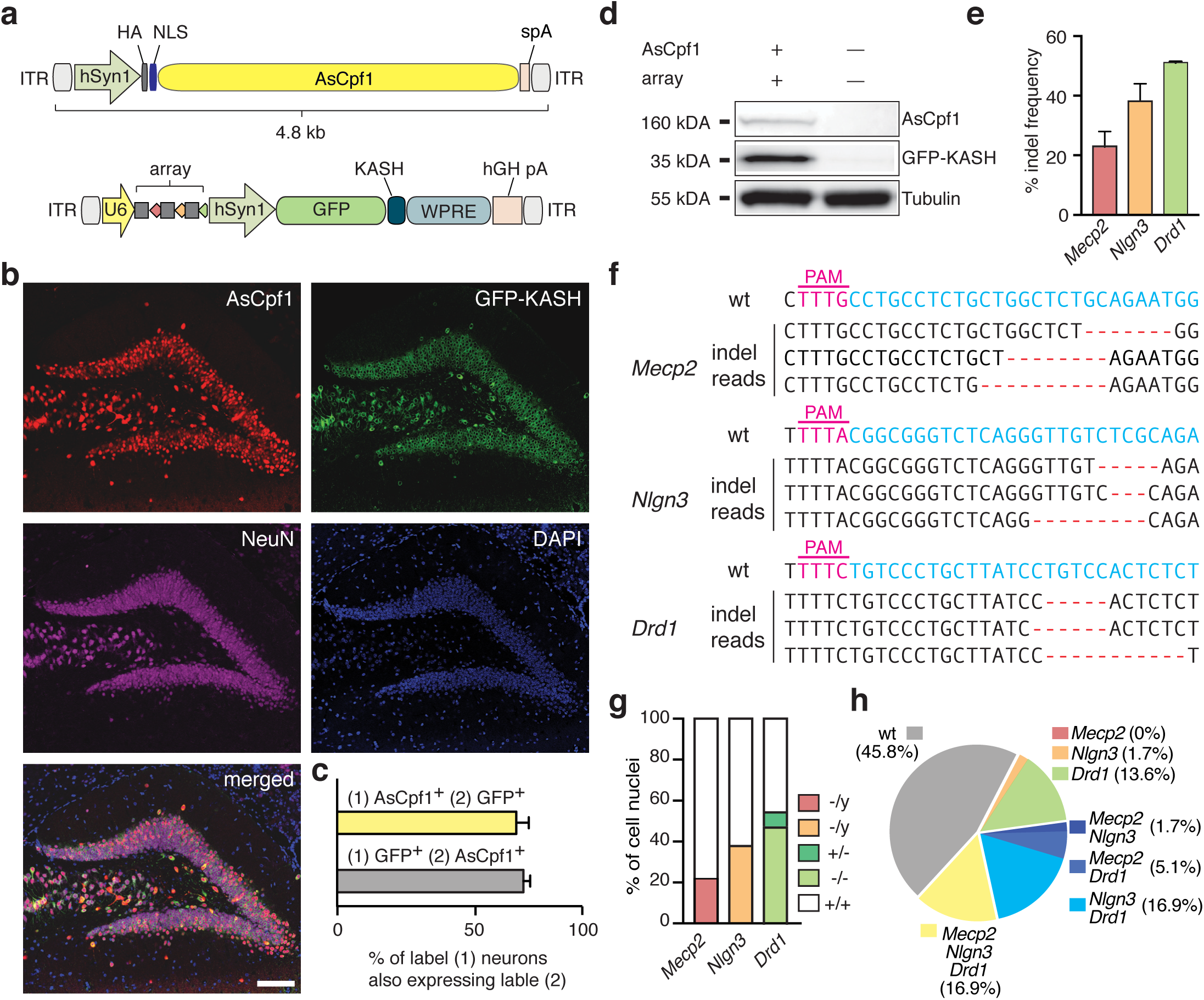
Delivery of AsCpf1 and multiplex gene editing in the mouse brain. (**a**) Schematic illustration of AAV vector design for multiplex gene editing. Bottom: grey rectangles, direct repeat; diamonds, spacer (red: *Mecp2*, orange: *Nlgn3*, green: *Drd1*) (**b**) Immunostaining of dorsal DG 4 weeks after stereotactic AAV injection (Representative image of *n* = 4 mice). Brain sections were co-stained with anti-HA (red), anti-GFP (green) and anti-NeuN (magenta) antibodies. Nuclei were labeled with DAPI (blue). Scale bar: 100 um. (**c**) Quantification of neurons efficiently transduced by the dual-vector system (*n* = 581 nuclei from 3 mice). (**d**) Western blot analysis of DG expressing HA-AsCpf1 and GFP-KASH (Representative blot from *n* = 4 mice). (**e**) NGS indel analysis of modified *Mecp2*, *Nlgn3* and *Drd1* loci in single nuclei (*n* = 59 cells from 2 male mice, error bars represent mean ± SEM). (**f**) Representative mutation patterns detected by NGS. Blue, wild-type (wt) sequence; red dashes, deleted bases; PAM sequence marked in magenta. (**g**) Fraction of mono-and biallelic modifications of autosomal gene *Drd1* is shown (*Mecp2* and *Nlgn3*: x-chromosomal). (**h**) Analysis of multiplexing efficiency in individual cells. ITR, inverted terminal repeat; HA, hemagglutinin tag; NLS, nuclear localization signal; spA, synthetic polyadenylation signal; U6, Pol III promoter; hSyn1, human synapsin 1 promoter; GFP, green fluorescent protein; KASH, Klarsicht, ANC1, Syne Homology nuclear transmembrane domain; hGH pA, human growth hormone polyadenylation signal; WPRE, Woodchuck Hepatitis virus posttranscriptional regulatory element.

Although multiplex gene editing is possible with Cas9, expression of several sgRNAs from one construct requires either multiple promoters (Kabadi et al., 2014; Sakuma et al., 2014), incorporation of long recognition sites for endogenous nucleases (Xie et al., 2015) or co-expression of a specialized nuclease cutting a small sequence between sgRNAs (Nissim et al., 2014; Tsai et al., 2014). This in turn requires either relatively large constructs, with cloning strategies not amenable for library cloning, or simultaneous delivery of multiple plasmids. Both are problematic for multiplex screens where viral delivery with low MOI is required as well as for *in vivo* applications using viral vectors with limited DNA packaging capacity.

In contrast, the ability of Cpf1 to process its own CRISPR array can be harnessed for multiplex genome editing both in vitro and in vivo using a single transcript driven by U6. Cpf1 only requires one Pol III promoter to drive several small crRNAs (39nt per crRNA). Hence, this system has the potential to simplify guide RNA delivery for many genome editing applications where targeting of multiple genes is desirable.

## Acknowledgements

We would like to thank F. A. Ran for helpful discussions and overall support, and Bas Cartigny and Jara van den Bogaerde for technical assistance; R. Macrae for critical reading of the manuscript, and the entire Zhang laboratory for support and advice. M.H was supported by the Human Frontiers Scientific Program. O.A.A. is supported by a Paul and Daisy Soros Fellowship and a Friends of the McGovern Institute Fellowship. J.S.G. is supported by a D.O.E. Computational Science Graduate Fellowship. E.D.G is supported by the National Institute of Biomedical Imaging and Bioengineering (NIBIB), of the National Institutes of Health (5T32EB1680). K.S. is supported by an NIH grant GM10407, Russian Science Foundation grant 14-14-00988, and Skoltech. J.v.d.O. is supported by Netherlands Organization for Scientific Research (NWO) through a TOP grant (714.015.001). F.Z. is supported by the NIH through NIMH (5DP1-MH100706 and 1R01-MH110049), the New York Stem Cell, Poitras, Simons, Paul G. Allen Family, and Vallee Foundations; and David R. Cheng, Tom Harriman, and B. Metcalfe. F.Z. is a New York Stem Cell Foundation Robertson Investigator. The authors plan to make the reagents widely available to the academic community through Addgene and to provide software tools via the Zhang lab website www.genome-engineering.org(www.genome-engineering.org).

